# Targeting PRMT5 Across Cancers and AI-Driven Discovery of Candidate Compounds

**DOI:** 10.64898/2026.10.19.760131

**Authors:** Sohel Rana, Md. Sanowar Hossan, Abdullah Al Rafi, Deri Damien Hagidok, Alok Kumer, Md. Enamul Kabir Talukder, Rasel A. Al-Amin

## Abstract

Protein arginine methyltransferase 5 (PRMT5) is a promising anticancer target, yet existing inhibitors shown limited clinical efficacy. Here, we paired multi-omics pan-cancer profiling with a machine learning virtual screening pipeline. Bioinformatics analyses with TIMER3, GEPIA2, UALCAN/CPTAC, the Human Protein Atlas, cBioPortal, STRING and Enrichr revealed widespread of PRMT5 mRNA expression, protein abundance, survival, genetic alteration, immune infiltration and co-expression networks with tumor-specific prognostic impact in GBM, LGG and KIRC. A deep neural network screened about 1.5 million PubChem compounds and deep neural network reached a ROC-AUC = 0.9857 and prioritized three candidates, CID 44345182, CID 2794188 and CID 44345334. Subsequent molecular docking against PRMT5 (PDB ID: 4X61), ADMET filtering and toxicity prediction, and 100 ns molecular dynamics (MD) simulations identified CID 44345182 as the potential hit, similar the docking scores of −9.8 compared with - 9.2 kcal/mol for the reference ligand in binding affinity and trajectory stability. These findings clarify the context-dependent roles of PRMT5 and further experiment validations are needed to establish CID 44345182 as a promising candidate.

## INTRODUCTION

Cancer remains a major global health challenges, highlighting the need for effective treatment strategies and better prevention [1]. Tumorigenesis involves genetic and epigenetic alterations that disrupt proliferation, survival, differentiation, and genome maintenance [2]. Protein arginine methylation is an important post-translational modification influencing transcription, RNA processing, DNA repair [3]. Aberrant PRMT5 expression and activity have been implicated in breast, lung, colorectal, prostate and haematological malignancies, supporting tumor proliferation, survival, invasion, and therapeutic resistance [4, 5]. A meta-analysis of 32 studies involving 15,583 patients associated elevated PRMT5 expression with advanced tumor stage and poorer overall and progression-free survival [6]. Mechanistically, PRMT5 can regulate cancer-associated RNA processing and DNA damage responses. For example, PRMT5-dependent splicing of DNA-repair genes contribute to chemoresistance in breast cancer stem cells, whereas disruption of PRMT5 mediated detained-intron processing and impair cancer-cell viability [7, 8]. These observations established a compelling rationale for targeting PRMT5.

Several small-molecule PRMT5 inhibitors have investigated into clinical evaluation. JNJ-64619178 demonstrated preclinical antitumor activity, while GSK3326595 highlighted challenges in balancing target inhibition with [9, 10]. A alternative homozygous MTAP deletion, reported in approximately 10-15% of cancers, which promotes methylthioadenosine (MTA) accumulation and vulnerability to PRMT5 inhibition [11]. MTA-cooperative inhibitors, including AMG 193, have shown preferential activity in MTAP-deleted solid tumors [12, 13]. Nevertheless, recent progress, tumor heterogeneity and therapeutic limitations motivate continued discovery of additional novel inhibitors. Natural products offer chemically diverse scaffolds for identifying promising anticancer Candidate compounds [14].

Computational screening can prioritize compounds before experiments testing, time and reducing discovery costs [15]. Machine-learning quantitative structure-activity relationships models learn from curated bioactivity data to large chemical libraries [16]. Property-matched decoys provide putative negative examples, while applicability-domain assessment helps identify predictions supported by the training space [17, 18]. Molecular docking and molecular dynamics (MD) simulations further evaluate binding plausibility and complex stability Previous PRMT5 discovery efforts include structure-based identification of T1551 for lung cancer candidate compound [19]. Emerging language-model agents can coordinate cheminformatics tools and workflows, although an agentic decision-support require clearly defined boundaries, handling model selection, applicability-domain checks, prediction aggregation and drug-likeness filtering without replacing the trained predictors [20, 21].

Here, we integrate pan-cancer characterization of PRMT5 with computational inhibitor discovery. Public datasets using TIMER, GEPIA2, UALCAN/ CPTAC and the Human Protein Atlas support analyses of transcript and protein expression, survival, genomic alteration (cBioPortal), stromal and immune infiltration interaction networks, and pathway enrichment (STRING, GEPIA2, Enrichr). PRMT5 bioactive records from ChEmBL (target CHEMBL1795116) together with 3,670 ChEMBL-derived PRMT5 actives and 3,235 property-matched decoys support comparative training of machine-learning and deep-learning classifiers built on RDKit descriptors and Morgan fingerprints **(Fig. 1)**. The selected model screen a PubChem library compounds, assisted by a bounded agentic layer for model selection, applicability-domain assessment, prediction aggregation, and drug-likeness filtering. Prioritized candidates undergo docking against PRMT5 structure 4X61, validation using EPZ 015666 as reference, ADMET and toxicity profiles, and MD simulations **(Fig. 1)**. Binding on integrated pan-cancer and machine-learning workflows recently applied for CDK2 and CDC7, this study aims to identify experimentally testable PRMT5 inhibitor candidates and clarify cancer context most relevant to therapeutic targeting [22, 23].

**Figure 1.**
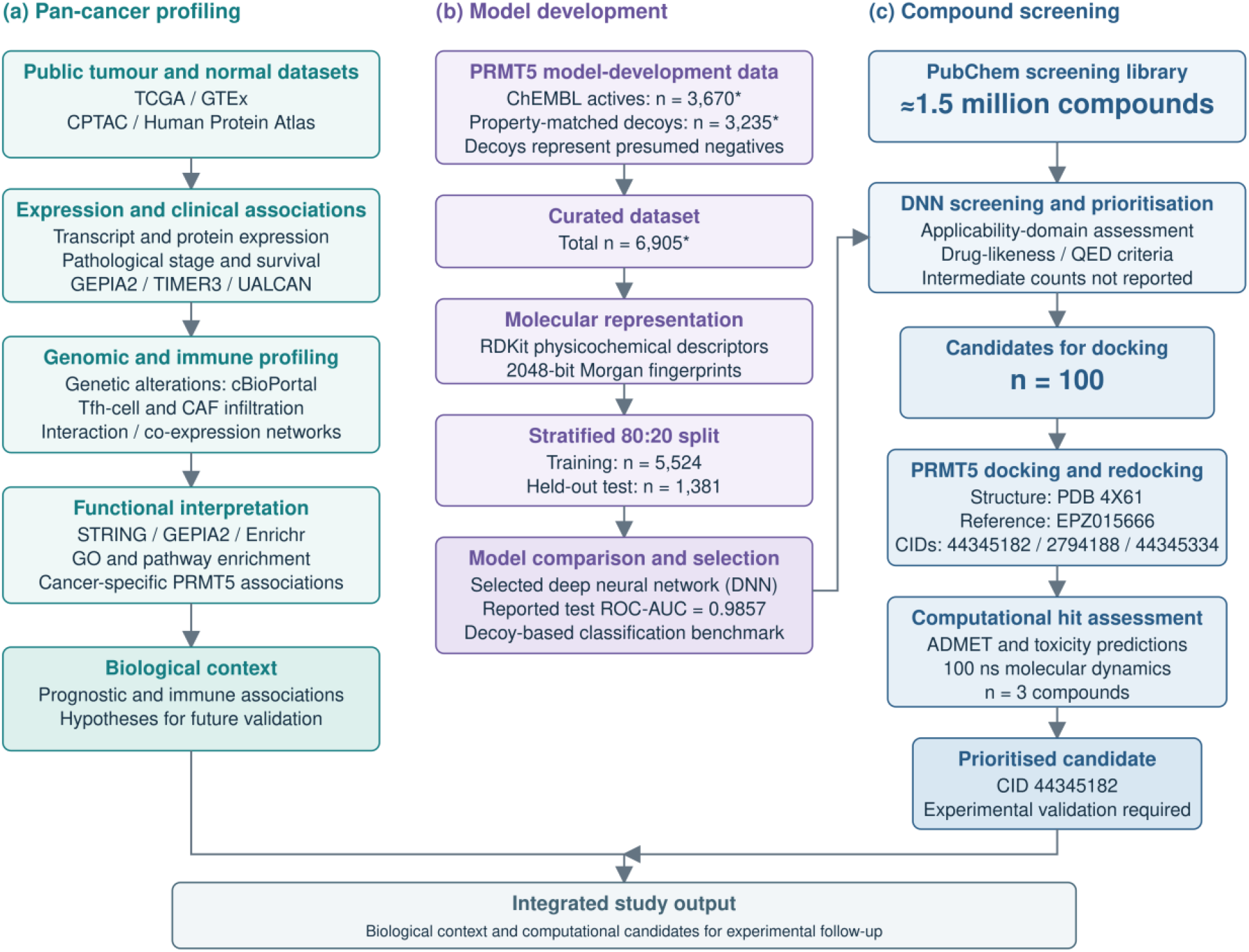
Study Design and Computational Compound-Screening Workflow. **(a)** Pan-cancer profiling of PRMT5 expression, clinical associations, genomic alterations and immune features, **(b)** Dataset preparation, molecular representation and deep neural network development. **(c)** Screening of approximately 1.5 million PubChem compounds, followed by docking, ADMET prediction and molecular dynamics simulations, prioritising CID 44345182 for experimental validation.

## METHODS

### Gene Expression, Proteomics Survival and Genetic Alteration Analysis

Transcriptional expression of PRMT5 across the Cancer Genome Atlas (TCGA) dataset or user-supplied data by employing six sophisticated algorithms, several cancer types and healthy tissues were analysed using TIMER3 (Tumor Immune Estimation Resource, version 3) database (http://timer.comp-genomics.org/) and Genotype-Tissue Expression (GTEx) database with GEPIA2 (http://gepia2.cancer-pku.cn/#analysis) [24, 25]. GEPIA2’s was utilised to examine gene expression levels “Expression Analysis-Stage Plot” generated violin plots across various pathological stages (I, II, III, and IV) using log2 (TPM+1) transformed data and ANOVA. Protein expression level was evaluated utilises data from the Clinical Proteomic Tumor Analysis Consortium (CPTAC) through UALCAN database (http://ualcan.path.uab.edu/analysis-prot.html), associated with the University of Alabama at Birmingham Cancer Data Analysis Portal, comprising 64 distinct cell lines, 48 healthy human tissues, and 20 tumor tissues [26, 27]. Immunohistochemistry (IHC) images of PRMT5 from the Human Protein Atlas (HPA) database, comprising 64 distinct cell lines, 48 healthy human tissues, and 20 tumor tissues [28]. Disease-free survival (DFS) and overall survival (OS) statistical data analysis were modelled via GEPIA2 using log-rank tests, with Cox Proportional Hazards (PH) Models for PRMT5 across all TCGA tumors [26]. Expression cutoffs of Low (50%) and High (50%) were established to delineate cohorts with elevated and diminished expression levels hazard ratios, and 95% confidence intervals (CI), depicted as a dotted line, were added to signify the accuracy of the hazard ratio (HR). PRMT5 mutations frequencies, structural mutated sites, copy number alteration (CAN), and genomic alteration impacts were mapped with cBioPortal (TCGA PanCancer Atlas dataset) which comprises 32 selected studies and a total of 10,967 samples (https://www.cbioportal.org/) [28]. Survival outcomes were compared between TCGA cancer cases with or without PRMT5 genetic alteration using Kaplan-Meier survival curves with log-rank tests.

### Immune Infiltration and Gene Enrichment Analysis

The TIMER3 Immune-Gene module was utilized to assess the Correlation between PRMT5 expression and T follicular helper (Tfh) cells were assessed using CIBERSORT, CIBERSORT-ABS, and ImmuCellAI algorithms, while cancer-associated fibroblasts (CAFs) were evaluated using EPIC (Extended Polydimensional Immunome Characterisation) algorithm. Associations between PRMT5 expression and immune-cell infiltration were quantify using Spearman’s rank correlations. Experimentally examined human protein PRMT5 interaction partners extracted from STRING database (https://string-db.org/) using “a minimum required interaction score” at “Low confidence (0.150)”, and “a maximum number of interactors displayed” in the first shell to (“no more than 50 interactors”). The top 100 co-expression genes were identify and selected corelations were examined using GEPIA2 utilised module “Expression Analysis-Similar Gene Detection” and TIMER2’s “Exploration-Gene_Corr” module. Pearson correlation analysis performed between PRMT5 and the selected genes, utilising log2-transformed TPM data for the dot plot visualisation and overlapping genes were identify using Jvenn (http://bioinformatics.psb.ugent.be/webtools/Venn/) [29]. Functional and pathway enrichment analysis were performed using Enrichr (https://maayanlab.cloud/Enrichr/) against Gene Ontology, KEGG, Reactome, and WikiPathways.

### Chemical Dataset Curation and Molecular Description

A set of 1.5 million bioactive compounds were retrieved from Pubchem (https://pubchem.ncbi.nlm.nih.gov/) [30]. A training dataset 3,670 experimentally validated active compounds (Chembl ID: CHEMBL1795116) was curated after removing duplicates and missing values via Python Pandas [31]. 3235 property-matched decoys molecules were generated following DUD-E benchmark as inactive compounds [32]. Dataset were spilt into an 80% training set of the total data, whereas, the remaining a 20% testing set kept for determining its performance using stratified sampling via Scikit-learn [31]. Physicochemical parameters including molecular weight (MW), lipophilicity (LogP), topological polar surface area (TPSA), hydrogen-bond counts, rotatable bonds, heavy atom count, ring count, and ring-related properties were calculated using RDKit-based cheminformatics workflow curated molecular dataset containing SMILES, while SMILES notation was specified for denoting the characteristics of molecules [33–36]. Compounds were categorized as active and inactive and Pearson correlation analysis were performed for the statistical visualization, and all analysis were performed by using Python libraries including RDKit, Matplotlib, Seaborn and Pandas **(Suppl. Fig. 6).** Tanimoto co-efficient similarity to training compounds was used to assess the applicability domain of external predictions based on molecular descriptor and molecular structure were additionally encoded using 2048-bit length Morgan fingerprints were computed using RDKit [33–40].

### Machine Learning Model & Framework

Multiple machine-learning algorithms were trained and benchmarked included Deep Neural Network (DNN), Convolutional Neural Network (CNN), Recurrent Neural Network with Gated Recurrent Unit (RNN-GRU), Gaussian Process (GP), Autoencoders + Logistic Regression (AE + LR), Bayesian Networks / Bayesian Classifier, Naïve Bayes (NB), k-Nearest Neighbours (k-NN), Decision Tree (DT), Principal Component Analysis + Logistic Regression (PCA + LR), Support Vector Machine (SVM), Artificial Neural Network–Multi-Layer Perceptron (ANN-MLP), Random Forest (RF), Random Forest Feature Selection (RF-FS), Gradient Boosting Machine (GBM), Molecular Fingerprints + Machine Learning Algorithms (FPs + ML), CatBoost, XGBoost, LightGBM and ebsemble pipelines [41]. Feature optimization was conducted using principal component analysis (PCA), Random-Forest-based feature selection, and autoencoder-based dimensionality reduction. Models performance was evaluated using ROC-AUC (primary matric/ indicator), through accuracy, balanced accuracy, weighted precision, weighted recall, weighted F1-score, and confusion matrices. All the models were ranked according to ROC-AUC, with weighted F1-score, and the best-performing model were selected for subsequent compound screening [36]. An Agentic AI decision framework was implemented to optimize model selection, compound prioritization, applicability-domain assessment, prediction aggregation, and drug-likeness filtering [21]. Applied QED scores, applicability-domain analysis, Lipinski-related drug-likeness properties filters [42].

### Protein Structure Preparation and Molecular Docking

The three-dimensional (3D) crystal structure of the target human protein PRMT5 (PDB 4X61) was retrieved from the RCSB Protein Data Bank (PDB) [43]. Heteroatoms, water molecules, and co-crystallized ligands were stripped using BIOVIA Discovery Studio 2021, followed by energy minimization in Swiss-PdbViewer (SPDBV version 4.1.0) [44–46]. Initial docking was performed using PyRx (version 0.8) [47]. The binding pocket of the target protein was identified using BIOVIA Discovery Studio 2021 with a receptor grid box centered at X = 22.881286, Y = - 47.593250, and Z = - 10.038750 [44]. High-affinity three top compounds (CID44345182, CID2794188, and CID44345334) were redocked using Schrödinger Maestro (version 11.1) with LigPrep and Protein Preparation Wizard, benchmarked against control (CID90241673).

### Pharmacokinetics (ADME), Toxicity Profiling and Molecular Dynamics (MD) Simulations

Pharmacokinetics behaviours, absorption, distribution, metabolism and excretion (ADME) properties were computed [48, 49]. In silico organ toxicity and toxicological assessment endpoints were predicted using the ProTox-II (https://tox-new.charite.De/) [49]. To validate protein-legend complexes stability, 100 ns period simulations were executed using the Desmond v6.3 module in Schrodinger (Release 2020-3) under Linux [50]. Complexes were solvated in a TIP3P water model within an orthorhombic periodic boundary box with a 10 Å buffer and neutralized [51]. Parameterization performed using the OPLS3e force field under the isothermal-isobaric condition, maintaining the temperature at 300 K and pressure at 1 atm [52]. Schrödinger Maestro version 9.5 was utilized to illustrate the MD simulation result. The simulation Trajectories were evaluated via the Simulation Interaction Diagram (SID) to track root mean square deviation (RMSD), root mean square fluctuation (RMSF), solvent accessible surface area (SASA), the radius of gyration (Rg), polar surface area (PSA) and Protein–ligand contacts (P-L contact) maintenance over experiment period.

## RESULTS

### Pan-cancer PRMT5 Expression and Proteomic Validation

Pan-cancer transcriptional profiling demonstrated significant PRMT5 overexpression across 15 various solid and haematological cancer types, including BRCA (Breast invasive carcinoma), CHOL (Cholangiocarcinoma), COAD (Colon adenocarcinoma), ESCA (Esophageal carcinoma), HNSC (Head and Neck squamous cell carcinoma), LIHC (Liver hepatocellular carcinoma), LUAD (Lung adenocarcinoma), LUSC (Lung squamous cell carcinoma), READ (Rectum adenocarcinoma), STAD (Stomach adenocarcinoma) (*P* < 0.001), BLCA (Bladder Urothelial Carcinoma) (*P* < 0.01), DLBC (Diffuse Large B-cell Lymphoma), THYM (Thymoma), Glioblastoma (GBM), and Low-Grade Glioma (LGG) (*P* < 0.05)-whereas significantly downregulated expression in KICH (Kidney Chromophobe) tumors (*P* <0.001), Pathological Stage correlation confirmed stage-dependent expression in in LIHC (Liver Hepatocellular Carcinoma); (*P* = 0.0289); KIRC (Kidney Renal Clear Cell Carcinoma) (*P* = 1.22e^-7^); and KICH (Kidney Chromophobe) (*P* = 0.0228) **(Suppl. Fig. 1c)**. Proteomic validation via UALCAN/CPTAC data and HPA immunohistochemistry images confirmed corresponding PRMT5 protein overexpression in Colon Cancer (CRC), Glioblastoma (GBM), Large Hepatocellular Carcinoma (LHCC), Lung Squamous Cell Carcinoma (LSCC), Lung Adenocarcinoma (LUAD), Pancreatic Ductal Adenocarcinoma (PDAC), Paediatric Brain Cancer (PDC), and Uterine Corpus Endometrial Carcinoma (UCEC), compared to normal samples, as illustrated in**(Suppl. Fig. 2a-b)**.

### Prognostic Relevance and Genomic Alternation

Survival analysis established high PRMT5 expression as a significant patient adverse prognosis indicator using TCGA data via GEPIA2 for overall survival (OS) outcomes in ACC (HR = 5.1, *p* = 0.00049), BLCA (HR = 1.4, *p* = 0.048), HNSC (HR = 1.3, *p* = 0.048), LAML (HR = 1.8, *p* = 0.036), LIHC (HR = 1.6, *p* = 0.0071), LUAD (HR = 1.3, *p* = 0.049), PAAD (HR = 1.7, *p* = 0.014), and SARC (HR = 1.6, *p* = 0.028), as well as reduced disease-free survival (DFS) in ACC (HR = 6.3, *p* = 7e-06), BLCA (HR = 1.5, *p* = 0.03), KICH (HR = 4.8, *p* = 0.047), LIHC (HR = 1.5, *p* = 0.0046), and PAAD (HR = 1.8, *p* = 0.0082) **(Suppl. Fig. 3)**. Conversely, low PRMT5 gene’s expression correlated with reduced overall survival (OS) for GBM (HR = 0.66, *p* = 0.022), KIRC (HR =0.69, *p* = 0.016), and LGG (HR = 0.59, *p* = 0.0043) and reduced disease-free survival (DFS) for GBM (HR = 0.61, *p* = 0.02), KIRC (HR =0.6, *p* = 0.0063), and LGG (HR = 0.66, *p* = 0.0087) **(Suppl. Fig. 3a-d)**.

Genetic alteration profiling identified an alteration frequency of PRMT5, over 3-4% in SKCM (Skin Cutaneous Melanoma) patients, attributed to “mutations” and “amplification”, in UCEC (Uterine Corpus Endometrial Carcinoma), the predominant forms of gene alteration were “mutation” and “amplification,” whereas in BLCA (Urothelial Bladder Carcinoma) and OV (Ovarian Serous Cystadenocarcinoma), alterations included “mutation,” “amplification,” and “deep deletion” of PRMT5 **(Suppl. Table 1** & **Fig. 2a-b)**. In total, mutation-level identified 105 PRMT5 variants predominantly comprising missense mutations; R534H/C mutation was specifically detected in cases of CRC (Colon Adenocarcinoma), MSTAD (Mucinous Stomach Adenocarcinoma), and UCEC (Uterine Endometrioid Carcinoma). These mutations result in missense variations in the PRMT5 gene, leading to substitutions of arginine (R) with histidine (H) or cysteine (C) at position 534 of the PRMT5 protein.

**Figure 2.**
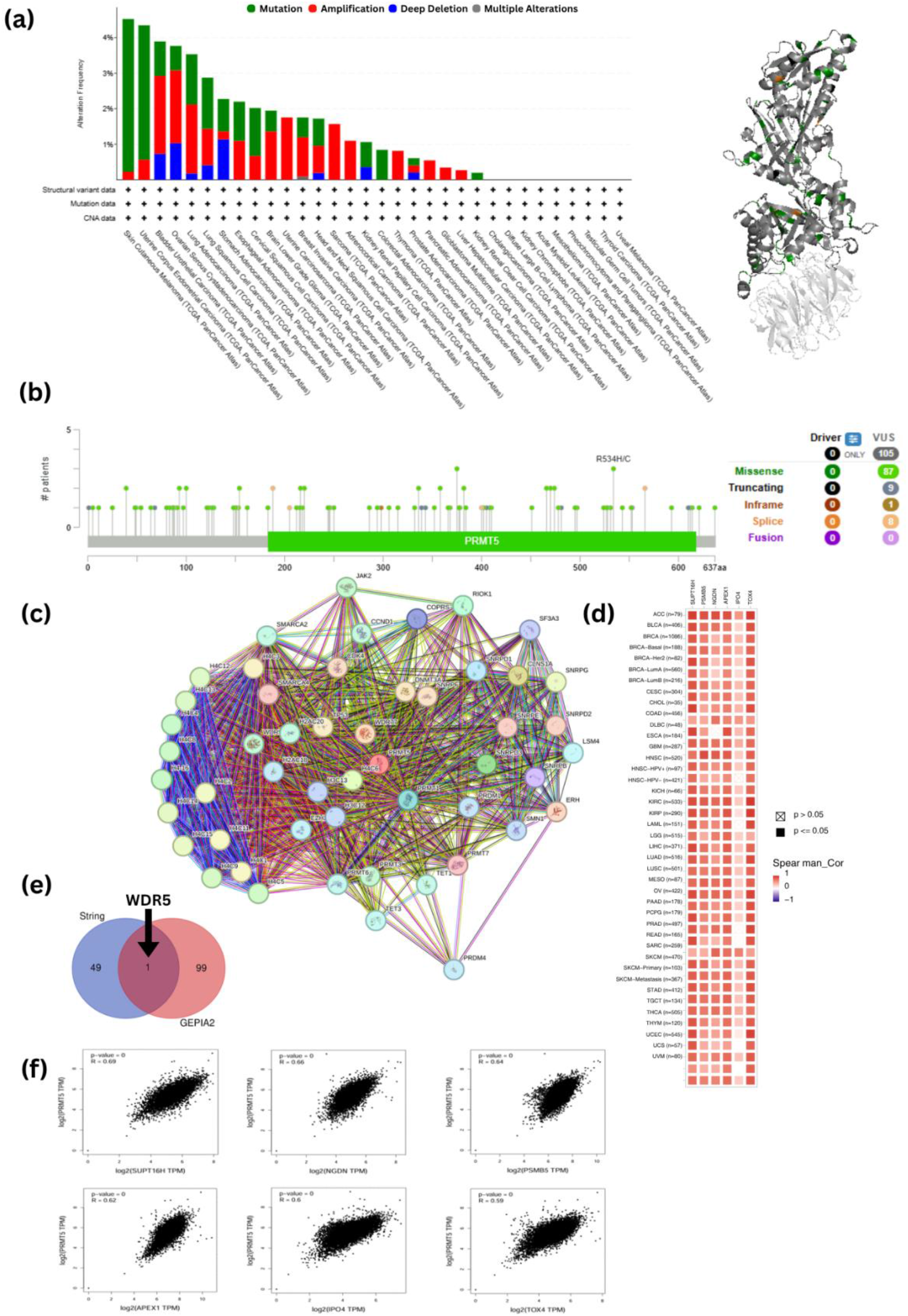
(a-b) Genomic alteration landscape and mutation mapping of PRMT5. Genetic variations landscape and mutation mapping of PRMT5 using cBioportal. **(a)** Frequency and types of PRMT5 genetic alterations and mutation across TCGA PanCancer Atlas studies derived from cBioPortal, alongside 3D structure mapping. **(b)** Domain distribution and specific mutation sites identified within the PRMT5 protein sequence. **(c-f) PRMT5 protein-protein interactome and co-expression network. (c)** STRING-derived functional protein interaction network displaying the top 50 PRMT5 interactors. **(d)** Co-expression correlation heatmap displaying positive expression relationships between PRMT5 and six top co-expressed genes (SUPT16H, NGDN, PSMB5, APEX1, IPO4, and TOX4). **(e)** Intersection of interaction and co-expression datasets identifying as a shared hub gene. **(f)** Individual correlation plots of PRMT5 expression against its top six co-expressed genes.

### Tumor Microenvironment Immune Infiltration & Splicing Interactome

EPIC algorithm mapping revealed strong positive cancer-specific correlation between PRMT5 expression and cancer-associated fibroblast (CAF) infiltration, significant positive correlations were observed in BRCA, BRCA-LumA, CESC, CHOL, ESCA, HNSC, LIHC, LUSC, PAAD, PRAD, SKCM, and THYM (*P* < 0.05). The strongest positive associations were observed in CHOL (*p* = 0.417), LIHC (*p* = 0.321), and THYM (*p* = 0.305), alongside universal significant negative associations were detected in LGG (*p* = - 0.127) and THCA (*p* = - 0.125) with T follicular helper cells (Tfh) [53, 54]. We found a negative correlation between PRMT5 expression and Tfh infiltration in BRCA, BRCA-LumA, HNSC, HNSC-HPV, KIRC, and LUAD cancers **(Suppl. Fig. 4)**.

Splicing interactome analysis identified 50 interactors utilizing the STRING program, top co-expressed genes, *SUPT16H* (R = 0.69), *NGDN* (R = 0.66), *PSMB5* (R = 0.64), *APEX1* (R = 0.62), *IPO4* (R = 0.60), and *TOX4* (R = 0.59) showed strong positive correlations with PRMT5 expression (all *P* < 0.001). The pan-cancer heatmap further confirmed positive correlations between PRMT5 and these six genes across multiple tumor types and identified a shared gene, interacting at WDR5 (**Fig. 2c-f)**. Functional enriched confirmed major involvement in mRNA-processing (*P* = 1.27 × 10^-16^, OR = 17.86), while GO Cellular Component showed strong enrichment in the nucleus (*P* = 1.03 × 10^-15^, OR = 5.16), and GO Molecular Function was primarily associated with mRNA binding (*P* = 5.21 × 10^-15^, OR = 14.35) **(Suppl. Fig. 5a-c)**. Reactome showed the strongest enrichment for RNA Metabolism (*P* = 5.24 × 10^-32^, OR = 18.51) and WikiPathways similarly identified mRNA Processing significantly enriched pathway (*P* = 1.85 × 10^-15^, OR = 29.02), KEGG identified the spliceosome as the most significantly enriched pathway (*P* = 3.35 × 10^-13^, OR = 22.55) **(Suppl. Fig. 5d-f)**.

### Deep Neural Network (DNN) QSAR Model Performance

The set of 3671 experimentally validated compounds (Chembl ID: CHEMBL1795116) were categorized as 1 active and 3234 decoys were categorized as 0 inactive. The whole dataset containing 6,905 molecular compounds which were categorised into training and testing subsets by using an partitioned into 80:20 ration approach using RDKit physicochemical characteristics, and Morgan circular fingerprints 2048-bit **(Suppl. Table 2 & Figure 3)**. A benchmark set of 3,670 active compounds (53.15%) and 3,235 decoy inactive compounds (46.85%). The independent test subset comprised 734 active compounds (53.15%) and 647 inactive compounds (46.85%), whereas the training subset included 2,936 active compounds (53.15%) and 2,588 inactive compounds (46.85%) **(Suppl. Table 3, Figure 3).** Among evaluated algorithms models, the Deep Neural Network (DNN) achieved superior predictive accuracy with a ROC-AUC value of 0.9857, effectively discriminating between active pRMT5 inhibitors from inactive decoy molecules. Tree-based ensemble models (Random Forest, XGBoost) also exhibited strong predictive capacity, confirming that circular fingerprints combined with global physical descriptors effectively capture nonlinear structure-activity relationships.

**Figure 3.**
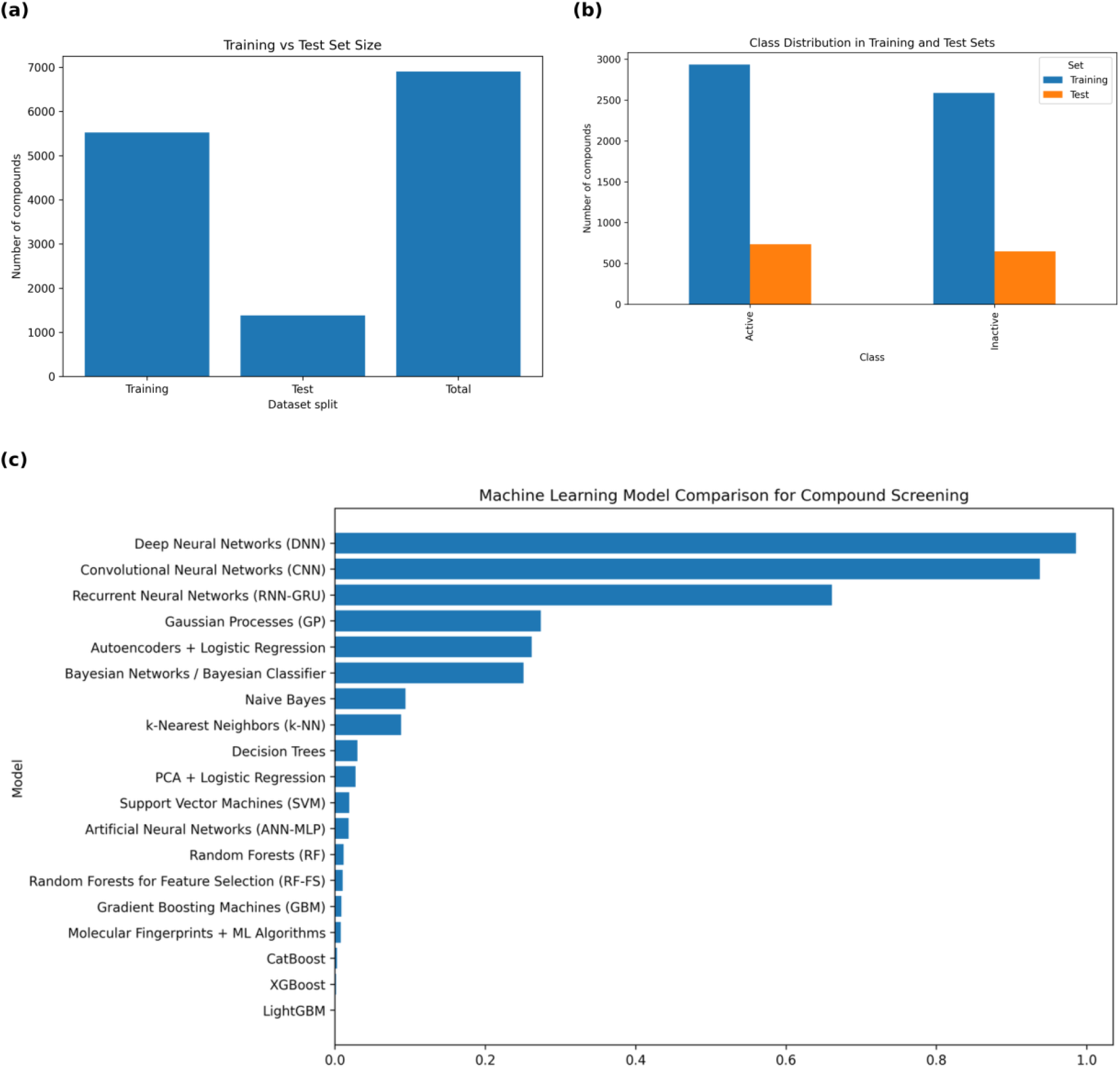
(a-b). Dataset partitioning for QSAR machine learning development. Summary of active and decoy compound distributions across training (optimization) and test (external validation) sets. **(c) Machine Learning Model Comparison for Compound Screening.** Among evaluated algorithms models, the Deep Neural Network (DNN) achieved superior predictive accuracy with a ROC-AUC value of 0.9857.

### Molecular Virtual and Active-Site Docking Analysis

Virtual screening of 100 phytochemical candidates’ compounds against the human PRMT5 target (PDB ID: 4X61) identified three primary Candidate compounds exhibiting high binding affinities; three candidates (IDs: CID 44345182, CID 2794188, and CID 44345334) with binding free energies (ΔG) of −9.8, −9.7, and −7.8 kcal mol⁻¹, respectively **(Fig. 4 & 5, Suppl. Table 4)**. All the three Candidate compounds demonstrated binding free energies comparable or superior to the experimental co-crystallized control ligand CID 90241673 (ΔG: 9.2 kcal mol⁻¹) **(Fig. 5 & Suppl. Table 4)**. Based on binding affinity, interactions with active site residues, ADME profiles, and toxicity predictions, three candidates (IDs: CID 44345182, CID 2794188, and CID 44345334), emerged as the most promising candidate compounds **(Table 1 & Fig. 4)**. Detailed active-site interaction profiling revealed specific binding pattern 3; CID 44345182 formed a conventional hydrogen bond with SER578, a carbon-hydrogen bond and a π-anion interaction with GLU444, a π-cation interaction with LYS333 and four aromatic interactions (2 π–π stacked and 2 π–π T-shaped) involving PHE327, PHE580, TYR324, TYR304, and a π-alkyl hydrophobic interaction with LEU A:319 **(Fig. 4a)**. CID 2794188 anchored via four hydrogen bonds with the residues ASP317, PRO314, LEU319, ASN394, and one Pi–Pi T-shaped interaction with residue TYR A:324 **(Fig. 4b)**. CID 44345334 engaged three hydrogen bonds with residues LYS517, ALA567, GLU534, and five Pi-Alkyl interactions with key hydrophobic residues (PHE492, LEU487, VAL495, VAL564, and LEU633) **(Fig. 4c)**. Observed strong binding stability at the PRMT5 active site, reinforcing its potential as a promising PRMT5 inhibitors (CID 44345182, CID 2794188, and CID 44345334) **(Fig. 4)**. Re-docking validation confirmed high conformational superimposition of the ligand binding modes accuracy, yielding a Root Square Deviation (RMSD) of 1.435Å between the re-docked pose and the co-crystallized native ligand (inhibitor CID 90241673). The re-docking procedure’s dependability was further confirmed by conformational superimposition of the ligand binding modes within the target protein **(Fig. 5)**.

**Figure 4.**
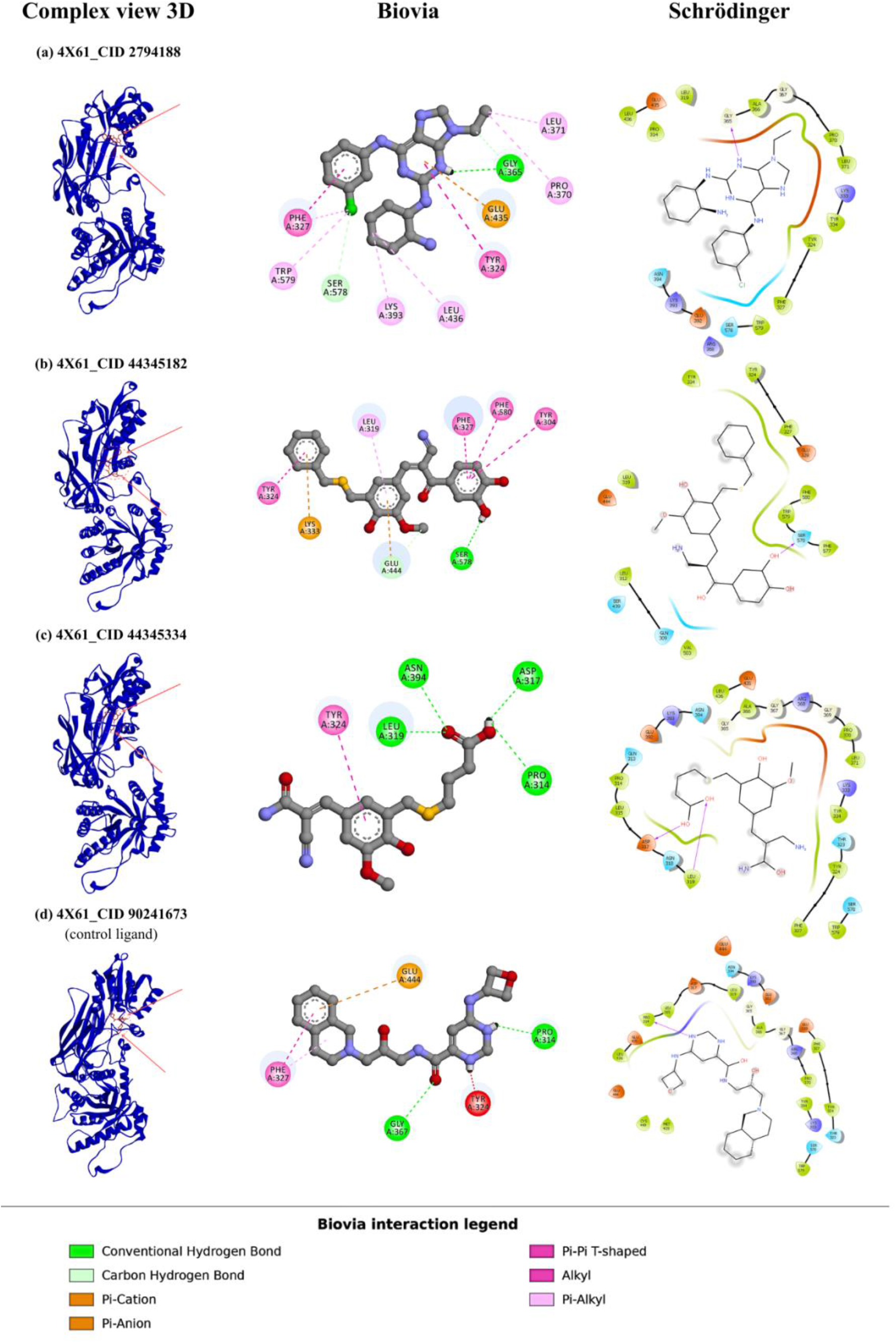
Predicted binding poses and interactions of candidate inhibitors with PRMT5 (PDB ID: 4X61). (a) CID 2794188, (b) CID 44345182, (c) CID 44345334, and (d) CID 90241673 (control ligand). Columns show 3D protein-ligand complexes view and interaction diagrams generated using BIOVIA and Schrödinger. Interaction types are colour-coded as indicated.

**Figure 5.**
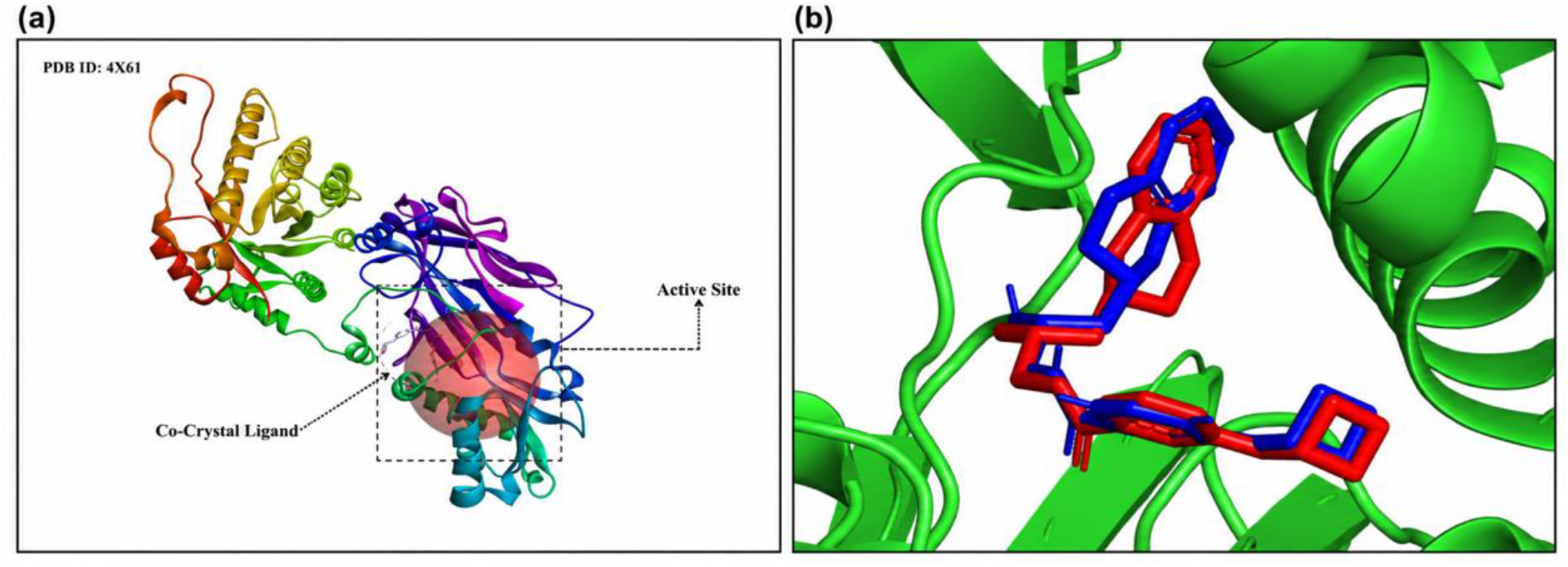
(a). Three-dimensional active-site architecture of human PRMT5. Three-dimensional Crystal structure of human PRMT5 (PDB ID: 4X61) bound to co-crystallized ligand (3XV), illustrating the catalytic binding pocket (red spherical region). **(b) Molecular re-docking pose superimposition.** Conformationally alignment of the re-docked co-crystallized ligand (CID 90241673) over its native crystallographic pose within the PRMT5 binding site, demonstrating RMSD of 1.435Å.

**Table 1.**
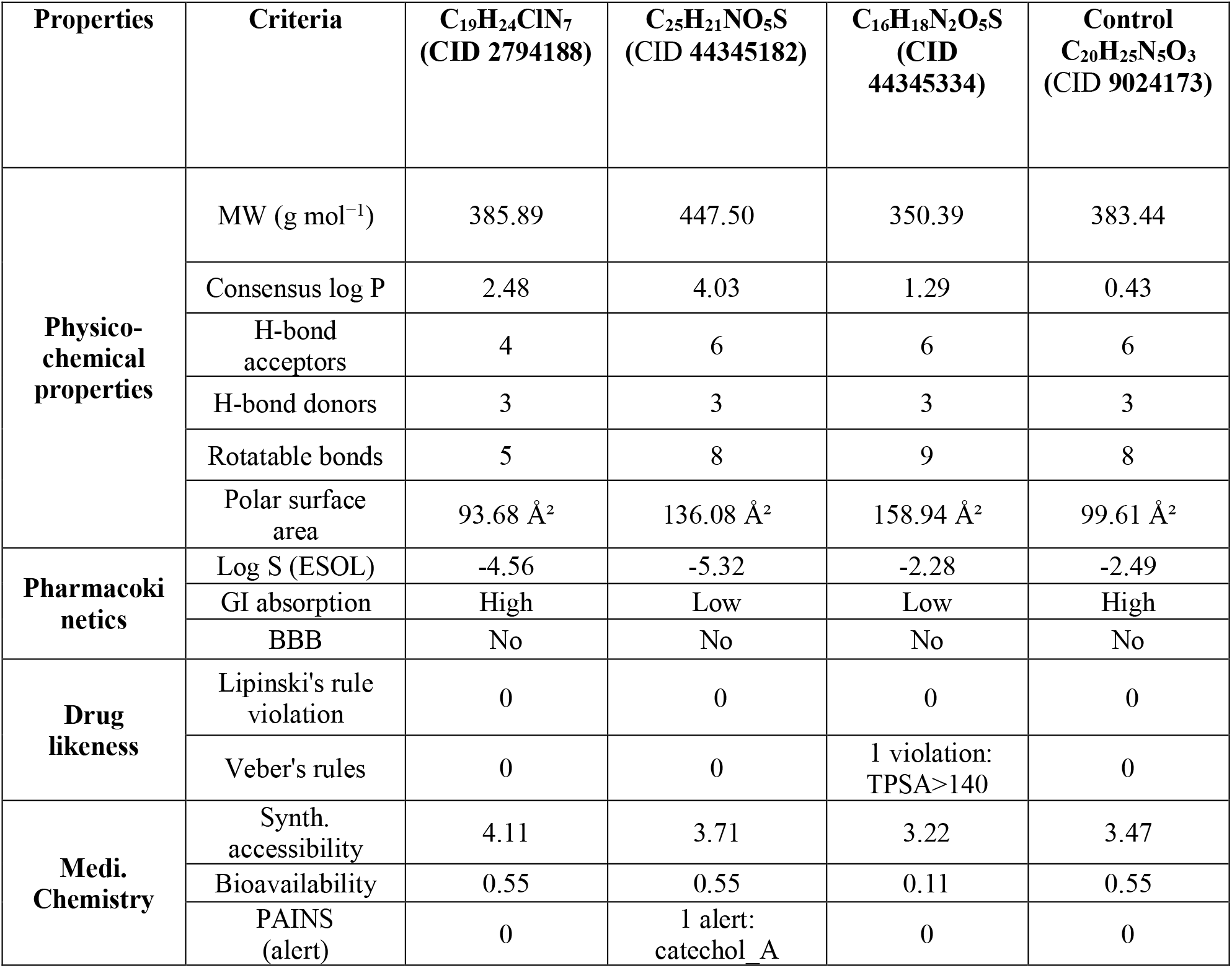
Physicochemical, Pharmacokinetic (ADME) and Bioavailability profiles of three candidate compounds along with the control.

The RMSD between the re-docked pose and the co-crystallized native ligand (inhibitor CID90241673) was 1.435Å, which is considerably below the universally accepted 2Å criterion, confirming the resilience of the docking procedure. This low divergence implies that the binding pose was accurately recreated, with the inhibitor occupying roughly identical locations and interacting with the same critical residues as the natural complex.

### Pharmacokinetic (ADME) and In Silico Toxicological Profiling

Evaluation via SwissADME (http://www.swissadme.ch/index.php) demonstrated all three candidate compounds (CID: 2794188, 44345182 and 44345334) adhere Lipiniski’s rule of five, number of rotatable bonds (nRotB) and polar surface area (PSA) without violation, with predicted the rule of five along with the log P values <5. The compounds are found to be moderate to high gastrointestinal (GI) absorption, greater synthetic accessibility, and clean PAINS (Pan-Assay Interference Compound) profiles indicating a low likelihood of assay interference or false positive result during biological screen. CID 44345182 and CID 44345334 were predicted to be non-permeable with the blood brain barriers (BBB), however, CID: 44345334 exhibited a lower oral bioavailability scored 0.11 indicating less suitable for oral dosage. Interference Compound) profiles indicating a low likelihood of assay interference or false positive result during biological screen. CID 44345182 and CID 44345334 were predicted to be non-permeable with the blood brain barriers (BBB), however, CID 44345334 exhibited a lower oral bioavailability scored 0.11 indicating less suitable for oral dosage. A comprehensive in silico toxicological analysis performed using ProTox-II (https://tox-new.charite.de/), confirmed clear safety profiles for compounds CID 2794188 and CID 44345182 across multiple toxicity end points including hepatotoxicity, neurotoxicity, cardiotoxicity, carcinogenicity, mutagenicity, cytotoxicity and predicted LD50, but CID 2794188 associated with possible toxicological risks **(Table 3)**. Compounds with lower LD₅₀ values indicated more toxic, whereas higher LD₅₀ values considered lower acute toxicity, a lower predicted acute median lethal dose CID 2794188 (160 mg/kg)> 44345182 and CID 44345334. The control ligand CID 9024173 also display higher toxicity profile risks associated with neurotoxic, central nervous system and blood brain barriers (BBB) permeability.

### Molecular Dynamic (MD) Simulation and Trajectory Analysis

#### Root Mean Square Deviation (RMSD) analysis

RMSD performed to evaluate the structural stability and conformational changes of the protein-ligand complex during the simulation. Lower RMSD fluctuations indicate a more stable protein backbone, while consistently low RMSD values provide favourable evidence for the structural stability of the docked complex [49]. The mean Cα RMSD values of the 4X61 complexes with CID 44345182 (3.337 Å), CID 2794188 (3.541 Å), and CID 44345334 (3.586 Å), and comparable to apo PRMT5 structure (3.221 Å) and control compound CID 90241673 (4.539 Å). The RMSD values of the candidate compound complexes remained comparable to that of the apo protein and lower than that of the control complex, indicating the maintenance of structural integrity and stable protein–ligand interactions throughout the simulation. Moreover, the overall RMSD profiles showed no significant structural deviations during the simulation period which supports the conformational stability of the protein and the persistent binding of the selected ligands within the active-site pocket **(Fig. 6)**.

**Figure 6.**
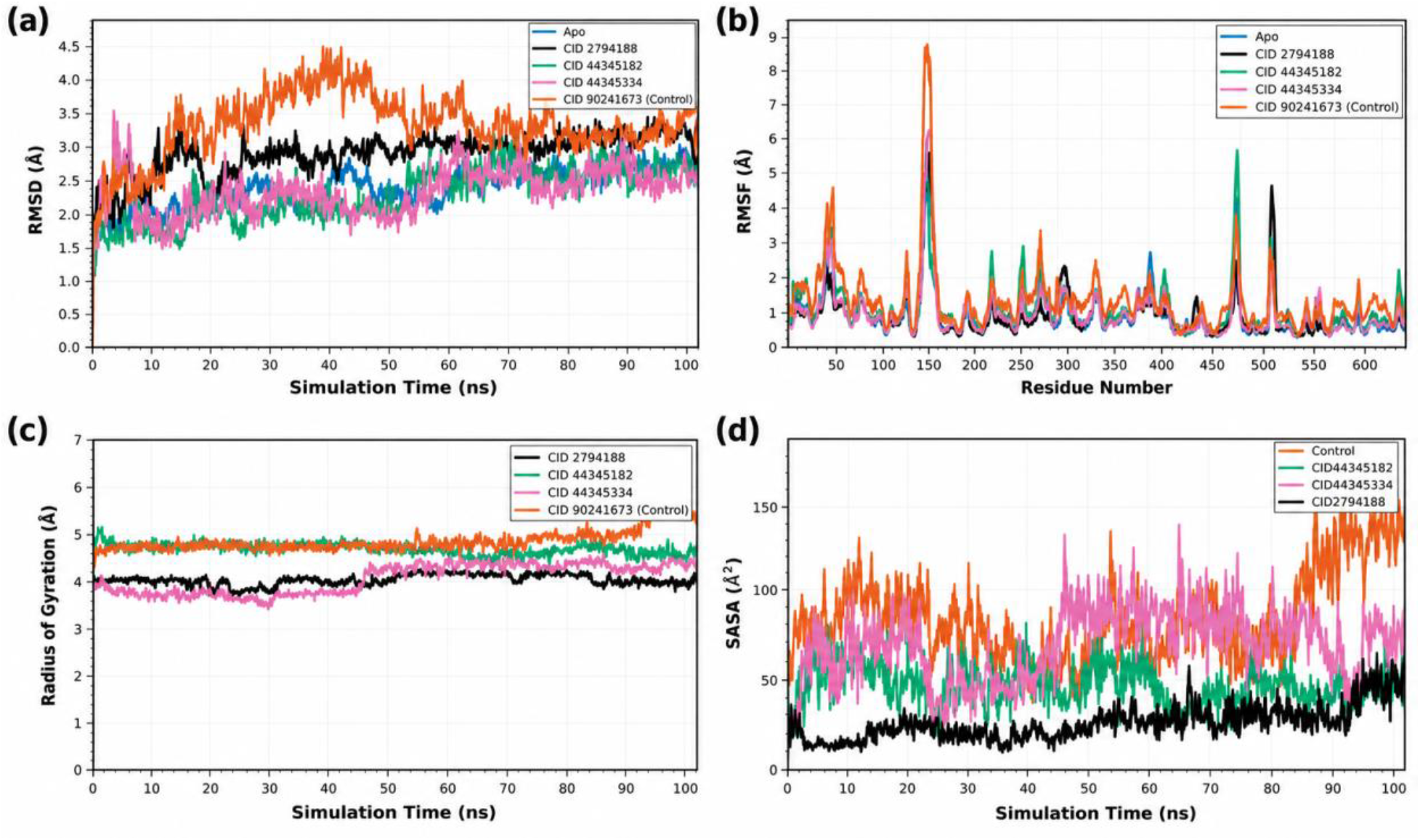
Molecular dynamics analysis of PRMT5–ligand complexes over 100 ns. **(a)** Cα Root-mean-square deviation (RMSD), **(b)** residue-wise root-mean-square fluctuation (RMSF), **(c)** radius of gyration (Rg), and **(d)** solvent accessible surface area (SASA) for the structural stability metrics over 100 ns for the three candidate compounds; CID 2794188 (black), CID 44345182 (green), CID 44345334 (pink), and control compound CID 90241673 (orange) with apo-PRMT5 (blue) included where indicated.

#### Root Mean Square Fluctuation (RMSF) analysis

RMSF demonstrated the average flexibility of individual amino acid residues in the target protein upon binding with the selected ligands, and measures the extent to which the atomic positions of protein residues fluctuate around their average positions over the simulation trajectory. RMSF analysis provide insights into the local structural flexibility and dynamic behaviour of the protein–ligand complexes [55]. RMSF analysis identified several regions primarily involved in ligand binding with relatively higher fluctuations, including with residue positions; PHE50 (38), ARG52 (40), LYS85 (73), ASN133 (121), VAL158 (146), ALA203 (191), LYS227 (215), VAL262 (250), TYR280 (268), LEU308 (296), GLN339 (327), PRO370 (358), PRO395 (383), TRP409 (397), PRO447 (435), SER478 (466), PRO515 (503), GLN561 (549), GLY595 (583), and ALA624 (612) (Fig. 5). Average RMSF values of 4X61 in complex were 1.16 Å (CID 44345182), 1.32 Å (CID 2794188), and 1.19 Å (CID 44345334), relative to 1.17 Å for apo form (PRMT5) and 1.62 Å for the control compound (CID 90241673) complex **(Fig. 6)**. RMSF values indicate fluctuations were restricted primarily to flexible loop regions associated with active-site dynamics, which supporting the structural stability of the selected protein–ligand complexes.

#### Radius of Gyration (Rg) Analysis

Rg reflects the compactness of a protein–ligand complex and describes how its atoms are spatially distributed around the molecular axis throughout the specified simulation period [53]. Changes in the radius of gyration further reflect the overall structural behaviour of the system, indicating either increased compactness or molecular expansion. Lower Rg values correspond to a more compact and structurally condensed configuration, mean Rg value were 4.92 Å (CID 44345182), 4.22 Å (CID 2794188), and 4.24 Å (CID 44345334), and 5.08 Å for the control compound (CID 90241673). Our finding indicate that binding of three candidate compounds does not induce substantial structural changes in the protein’s active site; maintained compactness of a protein–ligand complex compare to the control compound.

#### Solvent-Accessible Surface Area (SASA) analysis

SASA analysis evaluate the solvent-accessible surface area of the protein reflects the accessibility of hydrophobic and hydrophilic residue to the surrounding solvent. Over 100ns simulation, the 4X61 protein exhibit distinct SASA profiles depending on the bound compound, with an increase in SASA generally indicating greater exposure of the protein surface and associated confirmational expansion, SASA averages were 20.16 Å² (CID 2794188), 42.67 Å²(CID 44345182), and 65.61 Å² (CID 44345334), compared to 79.97 Å² for the control, indicating stable active-site condensation and spatial compactness throughout the simulation.

## DISCUSSION

PRMT5 serves as the primary type II methyltransferase responsible for catalysing monomethylation and symmetric dimethylation (SDMA) of arginine residues on histone and non-histone proteins using S-adenosylmethionine (SAM) [54]. Through these epigenetic and post-translational modifications, PRMT5 orchestrates critical physiological and cellular processes, including pre-mRNA splicing, DNA damage repair, ribosome biogenesis, metabolic regulation, and Golgi apparatus structure maintenance [56–60]. While aberrant PRMT5 expressed is frequently associated with tumor cell proliferation, survival, invasion, and therapeutic resistance across malignancies such as lung cancer, colorectal cancer, glioblastoma, clinical evaluation of first-generation several selective PRMT5 inhibitors (e.g., GSK3326595 and JNJ-64619178) has revealed limited monotherapy clinical efficacy and therapeutic window remains constrains, highlighting the need for improved PRMT5-targeting agents **(Table 2 & 3)** [5, 61–65].

**Table 2.** Physicochemical, pharmacokinetic (ADME) and drug likeness comparison of most potential candidate compound (CID 2794188) among with two others previously clinically evaluated of first-generation selective PRMT5 inhibitors (GSK 3326595 & JNJ 64619178).

| Properties | Criteria | C<sub>25</sub>H<sub>21</sub>NO<sub>5</sub>S<br>(CID 44345182) | Pemrametostat<br>(GSK 3326595) | Onametostat<br>(JNJ 64619178) |
| --- | --- | --- | --- | --- |
| Physico-chemical properties | MW (g mol<sup>-1</sup>) | 447.50 | 452.55 | 483.36 |
|  | Consensus log P | 4.03 | 1.33 | 2.08 |
|  | H-bond acceptors | 6 | 6 | 5 |
|  | H-bond donors | 3 | 3 | 4 |
|  | Rotatable bonds | 8 | 9 | 4 |
|  | Polar surface area ( Å<sup>2</sup>) | 136.08 | 110.69 | 136.10 |
| Pharmacokinetics | Log S (ESOL) | -5.32 | -3.03 | -4.48 |
|  | GI absorption | Low | High | Low |
|  | BBB | No | No | No |
| Drug likeness | Lipinski's rule violation | 0 | 0 | 0 |
|  | Veber's rules violation | 0 | 0 | 0 |
| Medi. Chemistry | Synth. accessibility | 3.71 | 3.90 | 4.16 |

**Table 3.** Predicted toxicity profiles including Hepatotoxicity, Neurotoxicity, Cardiotoxicity, Mutagenicity and LD_50_ of candidate compounds along with the control, and clinically evaluated of first-generation two selective PRMT5 inhibitors (GSK 3326595 & JNJ 64619178).

| **Toxicity Classification** | **CID 2794188** | **CID 44345182** | **CID 44345334** | **Control CID 9024173** | **GSK 3326595** | **JNJ 64619178** |
| --- | --- | --- | --- | --- | --- | --- |
| **Hepatotoxicity** | **Inactive** | **Inactive** | **Inactive** | **Inactive** | **Inactive** | **Inactive** |
| **Neurotoxicity** | Active | **Inactive** | **Inactive** | Active | Active | Active |
| **Cardiotoxicity** | **Inactive** | **Inactive** | **Inactive** | **Inactive** | **Inactive** | **Inactive** |
| **Carcinogenicity** | **Inactive** | **Inactive** | **Inactive** | **Inactive** | **Inactive** | Active |
| **Mutagenicity** | Active | **Inactive** | **Inactive** | **Inactive** | **Inactive** | Active |
| **Cytotoxicity** | **Inactive** | **Inactive** | **Inactive** | **Inactive** | **Inactive** | **Inactive** |
| **Predicted LD<sub>50</sub> (mg/kg)** | 160 | **1000** | **1000** | 200 | 200 | 57 |
| **BBB** | Active | **Inactive** | **Inactive** | Active | **Inactive** | Active |

Our multi-omics analysis demonstrates widespread but heterogenous dysregulation of PRMt5 at both the transcriptomic and proteomic levels. Rather than functioning as a uniform oncogene, the prognostic impact of PRMT5 is strictly context-dependent. Elevated PRMT5 expression represents a strong adverse prognostic factor in adrenocortical carcinoma (ACC; OS HR = 5.1, DFS HR = 6.3), pancreatic adenocarcinoma (PAAD), and bladder urothelial carcinoma (BLCA)2.

Conversely, lower PRMT5 expression correlates with reduced survival in glioblastoma (GBM), lower-grade glioma (LGG), and kidney renal clear cell carcinoma (KIRC) [66–71]. This bidirectional relationship highlights that transcript abundance does not inherently equate to functional dependency, as lineage-specific epigenetic landscapes, alternative splicing programs, and cellular heterogeneity dictate whether PRMT5 activity sustains survival or secondary tumor-suppressive networks.

Genomic profiling across 32 TCGA PanCancer Atlas cohorts revealed that direct genetic alterations in PRMT5 are rare, occurring less than 5% of cases even in the most altered malignancy (SKCM cutaneous melanoma, around 4.3% mutation). The identified 105 mutations consisted primarily variants of uncertain significance without recurrent driver mutations or genomic fusions. These findings indicate that genomic amplification or direct mutation of PRMT5 is not the predominant mechanism underlying PRMT5 dependency in human cancers These findings indicate that genomic amplification or direct mutation of PRMT5 locus is not the predominant mechanism underlying PRMT5 dependency in human cancers [72]. Instead, PRMT5 dependency is largely driven by collateral synthetic lethality, most notably following MTAP deletion. Homozygous deletion of MTAP candidates to the accumulation of 5′-methylthioadenosine (MTA), which selectively inhibits PRMT5 catalytic activity and renders MTAP-deficient cancer cells vulnerable to further PRMT5 blockade [73]. Consequently, clinical biomarker strategies should prioritize metabolic and genomic context (such as MTAP status) over PRMT5 transcript levels alone. Nevertheless, recurrent point mutations identified in this study, such as the R534H/C substitutions in colorectal and gastric carcinomas, warrant downstream structural characterization to evaluate their impact on catalytic efficiency, substrate selectivity, and inhibitor sensitivity.

Extending beyond tumor-cell-intrinsic mechanisms, our microenvironmental profiling demonstrates significant correlations between PRMT5 expression and specific stromal and immune components. PRMT5 expression correlated positive cancer-associated fibroblast (CAF) infiltration across cholangiocarcinoma (CHOL), LIHC and thymom (THYM), while displaying inverse correlations with T follicular helper (Tfh) cells in breasts, head and neck, and lung adenocarcinomas [70, 74]. PRMT5 suppresses antitumor immunity by by methylating key signalling intermediates within the cGAS-STING pathway and downregulating NLRC5 pathway-mediated MHC class I antigen presentation. Pharmacological inhibition PRMT5 has been shown interferon immune signalling and enhance efficacy of immune-checkpoint therapy [75]. However, because PRMT5 also regulated physiological lymphocyte maturation and activation, systemic inhibition may elicit opposing cell-type-specific immune responses. These findings provide a rationale for exploring combination strategies pairing PRMT5 inhibitors with immunotherapies in stroma-rich or immune-excluded tumor microenvironments.

Functional enrichment of the PRMT5 co-expression revealed robust convergence on pre-mRNA processing, RNA spliceosomal assembly, RNA transport, and mRNA surveillance across Gene Ontology, KEGG, Reactome, and WikiPathways databases. Splicing regulation represents the most well-validated biological dependency of PRMT5; symmetric dimethylation of Sm proteins is essential for snRNP biogenesis, and PRMT5 inhibition induces intron retention and exon skipping that preferentially destabilize transcripts required for cell cycle progression [76]. Secondary enrichment was observed in DNA damage repair, cell cycle control, proteasomal degradation, and nucleocytoplasmic transport pathways [77]. Mechanistically, these pathways intersect with splicing regulation, as aberrant pre-mRNA splicing directly alters the isoform composition of DNA repair factors and cell cycle regulators [77–79]. Together, these results suggest that PRMT5 may connect RNA-processing machinery with proliferative, genome-maintenance, and immune-regulatory programs rather than operating through a single isolated oncogenic pathway.

Cross-referencing the PRMT5 protein-interaction network and the co-expression dataset identified a single shared gene, WDR5. WDR5 is an essential component of histone methyltransferase complexes involved in chromatin remodelling, suggesting potential coordination between PRMT5-mediated arginine methylation and WDR5-dependent transcriptional control [80, 81]. From a translational perspective, the present findings support PRMT5 as a therapeutically relevant but biomarker-dependent cancer target. Early approaches to global PRMT5 inhibition established pharmacological tractability but also highlighted limitations related to therapeutic window and effects on normal PRMT5-dependent tissues [82]. More recent strategies exploit tumor-specific molecular contexts, particularly the MTA-PRMT5 dependency created by MTAP deletion, providing a stronger framework for precision targeting of PRMT5 [83]. Accordingly, expression alone is unlikely to be sufficient for patient selection; integration of PRMT5 expression with genomic background, metabolic state, RNA-processing dependency, and immune context may provide a more informative strategy for identifying responsive tumors.

To identify novel small-molecule inhibitors targeting PRMT5, we developed a machine learning framework trained on 3,670 active compounds and 3,235 property-matched decoy inactive. Feature selection combining 2048-bit Morgan circular fingerprints with global physical descriptors enabled the models to capture non-linear structure-activity relationship. Among the benchmarked architectures, the Deep Neural Network (DNN) model achieved superior classification performance on the independent test set (ROC-AUC = 0.9857). Although performance metrics on decoy-based datasets can overestimate accuracy compared to true experimental inactive, the classifier provided an efficient filter to prioritize large chemical libraries prior to docking. Deploying the DNN model alongside applicability-domain validation and drug-likeness filters successfully narrowed a 1.5 million PubChem compound library down to 100 candidate molecules. These candidates were then subjected to molecular docking, which is a widely used computational approach for evaluating the binding affinity and interaction patterns between small molecules and their target macromolecules. These candidates were then subjected to molecular docking, which is a widely used computational approach for evaluating the binding affinity and interaction patterns between small molecules and their target macromolecules.

In silico ADMET profiling confirmed that all three hit compounds adhere to Lipinski’s rule of five. CID 2794188 and CID 44345182 met Veber’s criteria, whereas CID 44345334 presented a single Veber violation due to elevated topological polar surface area (TPSA > 140 Å²) **(Table 1)**. Although SwissADME zero blood–brain barrier (BBB) for all candidate compounds, ProTox-II flagged CID 2794188 as BBB-permeable, neurotoxic, and mutagenic with a lower predicted median lethal dose (LD_50_ = 160 mg/kg), making it the least attractive candidate **(Table 3)**. In contrast, CID 44345182 and CID 44345334 showed clean predicted toxicity profiles (LD_50_ = 1000 mg/kg). Whereas CID 2794188 was predicted to be neurotoxic **(Table 3)**. Nevertheless, specific liabilities-such as CID 44345182 showed low predicted gastrointestinal absorption and a catechol PAINS alert, and CID 44345334 a low bioavailability score (0.11), - highlight the necessity for downstream chemical structural optimization **(Table 1)**. Over100 ns molecular dynamics (MD) simulation, all the three identified compounds complexes demonstrated structural stability and active-site retention. Mean backbone RMSD values (3.34 - 3.59 Å), and local atomic fluctuations (RMSF = 1.16-1.32 Å), closely mirrored the apo-PRTM5 structure (RMSD = 3.22 Å, RMSF = 1.17 Å) and remained substantially lower than the control the control complex (RMSD = 4.54 Å, RMSF = 1.62 Å). Furthermore, Radius of Gyration (Rg) and Solvent Accessible Surface Area (SASA) profiles were consistently lower across the candidate compound complexes than in the control complex, indicating sustained spatial compactness and reduced solvent exposure throughout the simulation. While these computational findings provide a prioritized chemical framework, experimental in vitro and in vivo validation remains required to confirm biological efficacy, selectivity, and safety.

## CONCLUSION

PRMT5 is broadly overexpressed across a diverse tumor type at both the transcript and protein levels, yet correlation with patient survival varies substantially across cancer type and lacks recurrent driver alterations. Functional co-expression and pathway enrichment analyses indicate that PRMT5 dependency is primarily centered on RNA processing and pre-mRNA splicing networks, along secondary involvement in DNA repair, cell-cycle regulation control, and immune pathways. Consequently, PRMT5 is best conceptualized as a context-dependent therapeutic target rather than a universal oncogenic driver. This study integrated pan-cancer bioinformatic profiling with deep neural network-guided virtual screening (ROC-AUC = 0.9857), molecular docking, ADMET filtering and toxicity prediction, and 100 ns molecular dynamics simulation to prioritize three candidates PRMT5 compounds: CID 44345182, CID 2794188, and CID 44345334. Among the evaluated hits, CID 44345182 demonstrated the most favorable overall profile, combining top-ranking binding affinity (−9.8 kcal mol⁻¹), a clean predicted toxicity profile, and stable dynamic trajectories. PRMT5 functions as a context-dependent therapeutic target predominantly linked to pre-mRNA splicing and RNA processing pathways. All findings rely on computational predictions; the deep neural network classifier (ROC-AUC = 0.9857) was trained against property-matched decoys rather than confirmed experimental inactive, MD trajectories were limited to 100 ns without binding free energy calculations, and no experimental wet-lab validation was conducted. Furthermore, database evaluations may be influenced by tumor purity and cohort bias, and analyses were not stratified by MTAP deletion status. Future work should evaluate these candidate compounds in biochemical and cell-based assays across MTAP-deficient models, assess selectivity across PRMT family members, execute thermodynamic binding free energy calculations, and perform candidate compound optimization.

## AUTHOR CONTRIBUTIONS

S.H., M.S.H. and R.A.A.-A.: study design, and investigation. M.S.H.: Pan-cancer profiling methodology, data curation and analysis, writing draft version, and review. S.H.: Development of the AI-driven methodology in consultation with R.A.A.-A., data curation and analysis, writing draft version, and review. A.A.R., D.D.H., and A.K.: Methodology, data analysis and curation, and writing initial draft version. M.E.K.T.: Molecular dynamics simulation analysis and data curation. R.A.A.-A.: Conceptualization the idea, resources, validation, supervision, re-writing original version, review and editing, and preparation of the final manuscript. All authors read and approved the final manuscript.

## CORRESPONDING AUTHOR

**\*Rasel A. Al-Amin** (orcid.org/0000-0002-0762-9034)

## ORIGINALITY DECLARATION

The work presented in this manuscript has not been published previously and is not under consideration for publication elsewhere, in whole or in part.

## ACKNOWLEDGEMENTS

Declaration of AI-assisted language editing: The authors may be used artificial intelligence tools solely to improve the grammar, language, and clarity of this manuscript. All AI-assisted revisions were reviewed and approved by the authors, who take full responsibility for the manuscript’s scientific content, data interpretation, and conclusions.

