## Supplementary Figures and Tables for "Targeting PRMT5 Across Cancers and AI-Driven Discovery of Candidate Compounds"

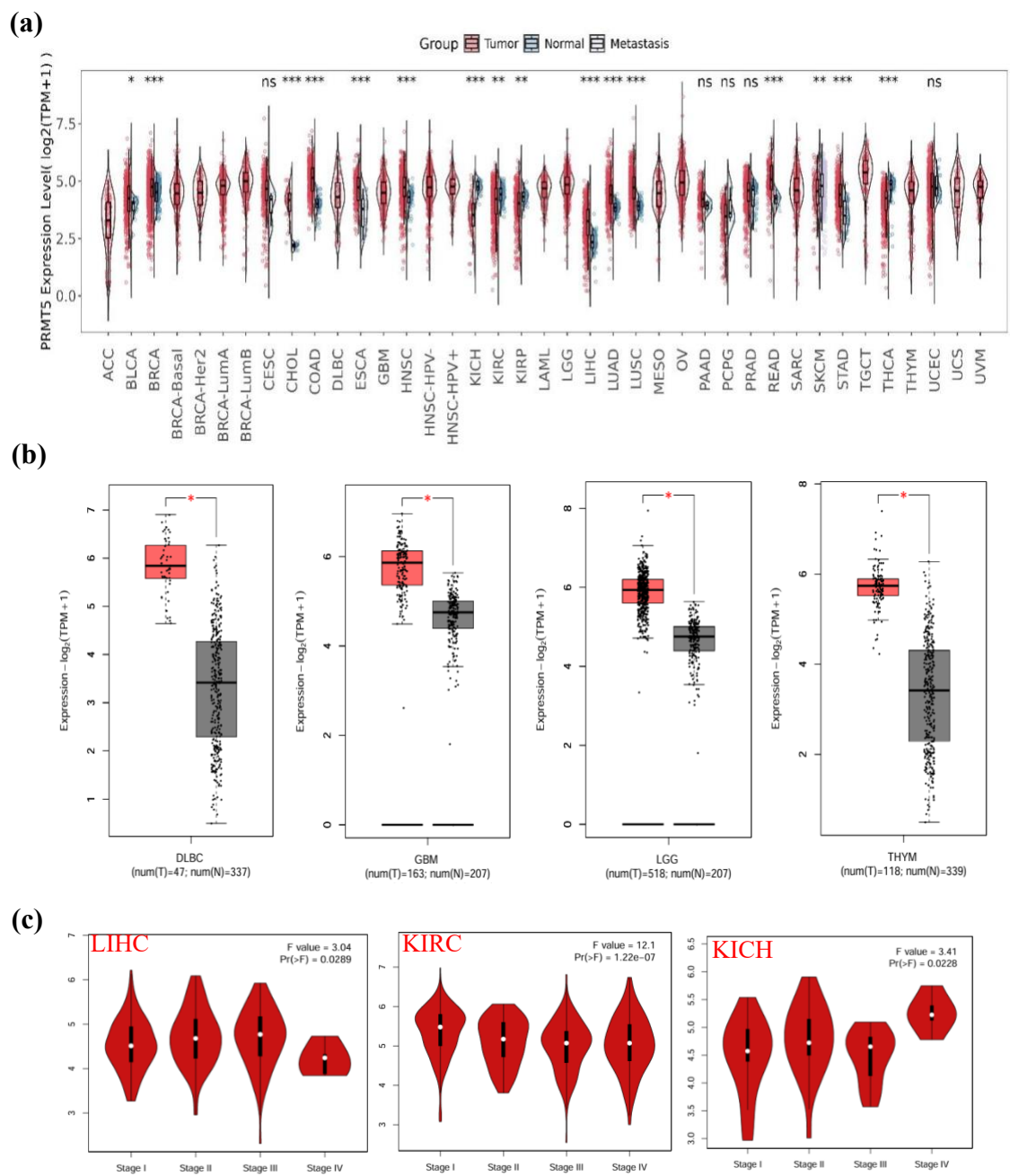

**Supplementary Figure 1. Pan-cancer transcriptomic expression and stage-dependent profiling of PRMT5.** **(a)** Comparative expression levels of PRMT5 across TCGA tumor types and matched normal tissues evaluated via TIMER2 ($*p < 0.05$, $**p < 0.01$, and $***p < 0.001$). **(b)** PRMT5 expression in DLBC, GBM, LGG, and THYM normal tissues from GTEx database served as controls displayed via box plots ($**p < 0.01$ indicating statistical significance). **(c)** Pathological stage-dependent PRMT5 gene expression across clinical phases (I, II, III, and IV) for LIHC, KIRC, and KICH derived from TCGA data.

(a)

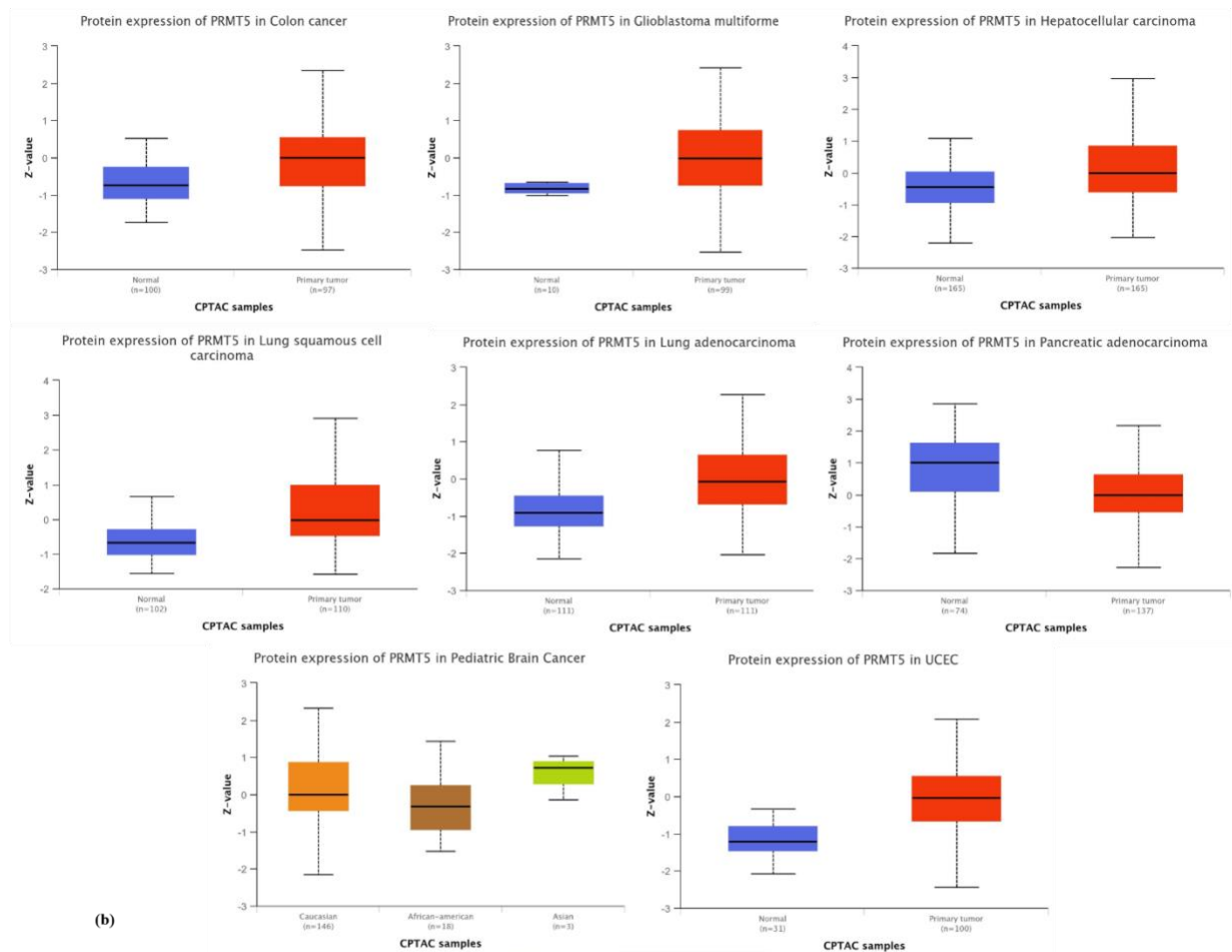

(b)

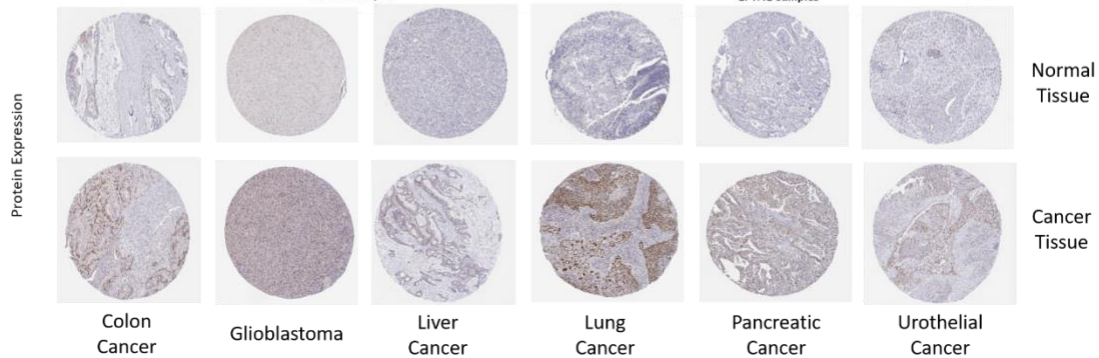

**Supplementary Figure 2.** Proteomic validation and tissue-level expression of PRMT5. **(a)** Total PRMT5 protein levels in primary tumor tissues versus normal controls evaluated using CPTAC dataset cohorts. **(b)** Representative immunohistochemical (IHC) staining profiles showing PRMT5 protein localization in normal and cancerous tissues from the Human Protein Atlas (HPA).

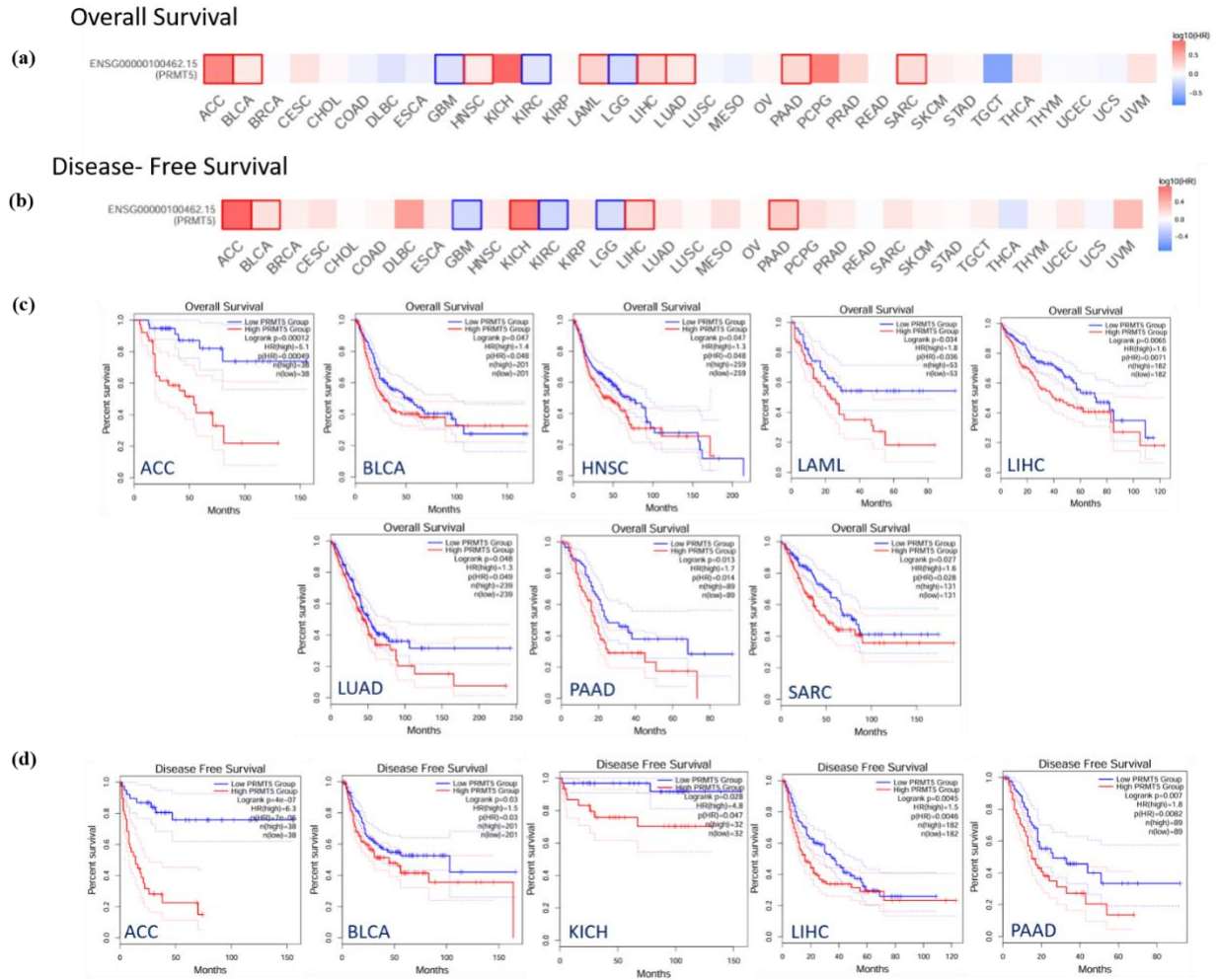

**Supplementary Figure 3. Prognostic significance of PRMT5 expression across human cancers.** Kaplan–Meier survival analysis conducted via GEPIA2 showing (a) overall survival (OS), (b) disease-free survival (DFS), and (c) OS curves for high versus low PRMT5 expression in ACC, BLCA, HNSC, LAML, LIHC, LUAD, PAAD, and SARC, (d) DFS curves in ACC, BLCA, KICH, LIHC, and PAAD, all results demonstrating statistical significance ($p < 0.05$).

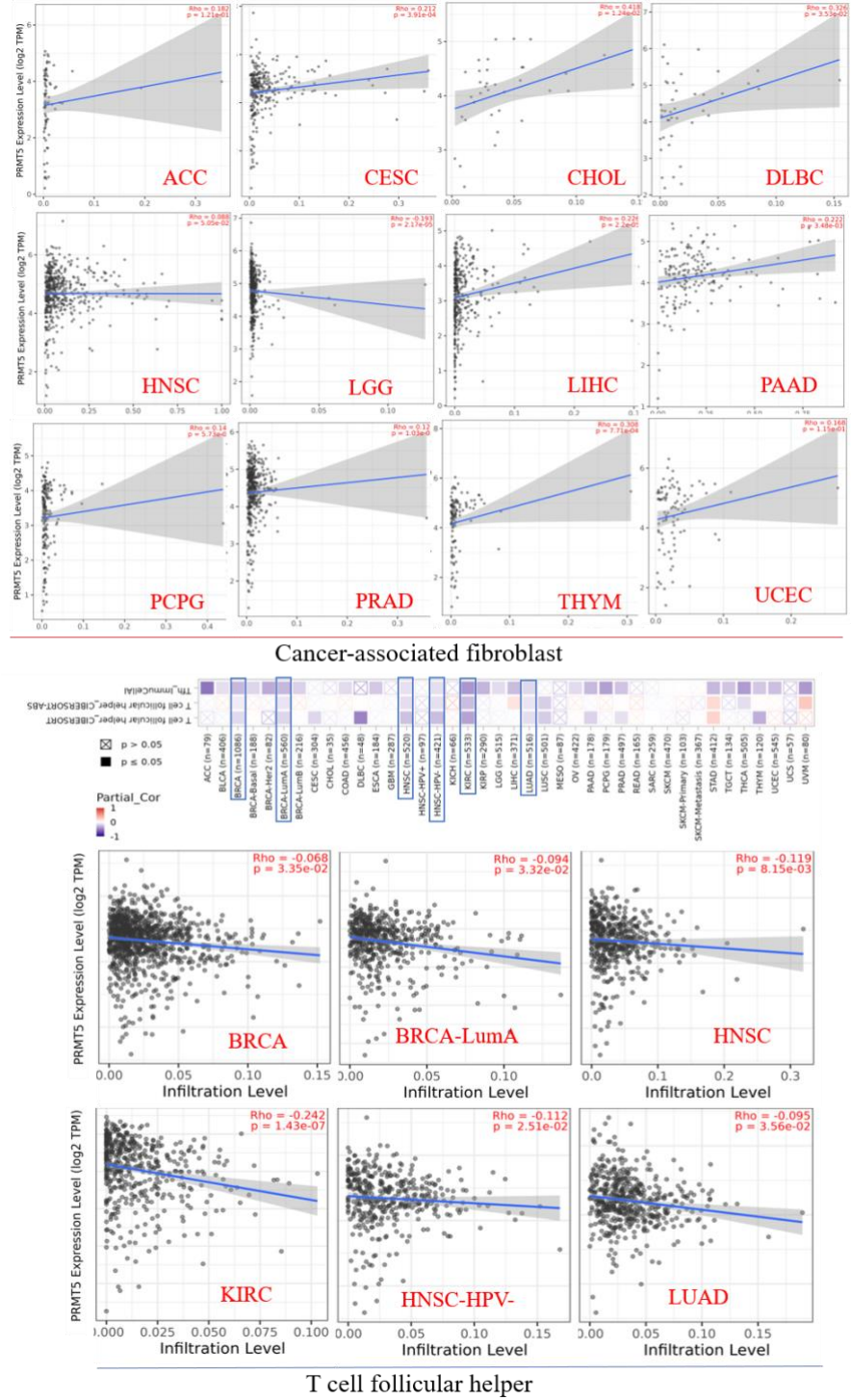

**Supplementary Figure 4. Immune cell infiltration analysis of PRMT5 across TCGA cohorts.** Correlation between PRMT5 transcript expression and microenvironmental infiltration of T follicular helper (Tfh) cells and cancer-associated fibroblasts (CAFs) estimated across algorithms. Correlation between PRMT5 transcript expression and immunological microenvironmental infiltration of T follicular helper (Tfh) cells and cancer-associated fibroblasts (CAFs) estimated across algorithms.

(a)

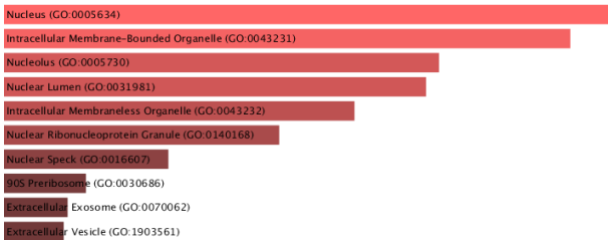

### GO BIOLOGICAL PROCESS

(b)

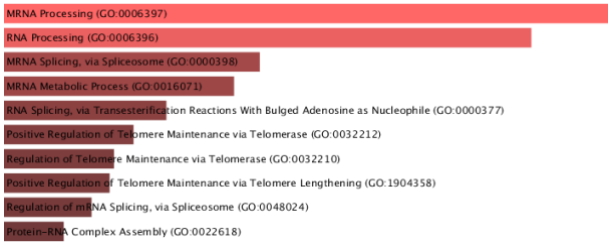

### GO CELLULAR COMPONENT

(c)

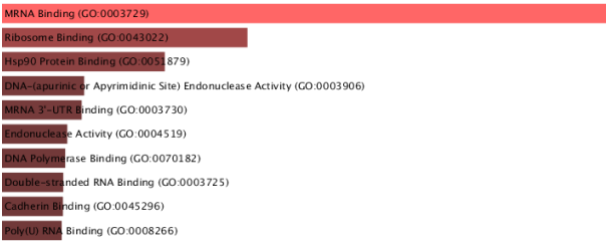

### GO MOLECULAR FUNCTION

(d)

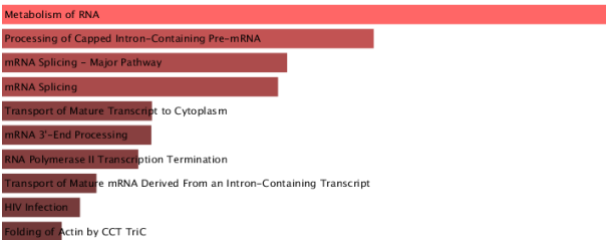

### REACTOME

(e)

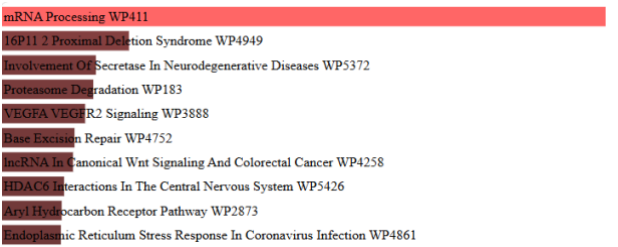

### WikiPathway

(f)

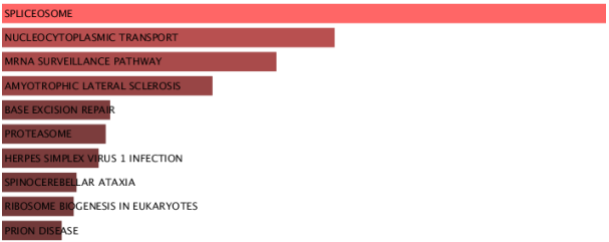

### KEGG

**Supplementary Figure 5. Functional pathway enrichment analysis of PRMT5-associated genes.** PRMT5 enrichment analysis, bar diagrams generated using Enrichr (https://maayanlab.cloud/Enrichr/) indicate a significant association between PRMT5 and different metabolic pathways, and a lighter colour indicates a more significant result. (a) GO Biological Process 2026, (b) GO Cellular Component 2026, (c) GO Molecular Function 2026, (d) Reactome Pathway 2024, (e) WikiPathways 2024 Human, and (f) KEGG 2026.

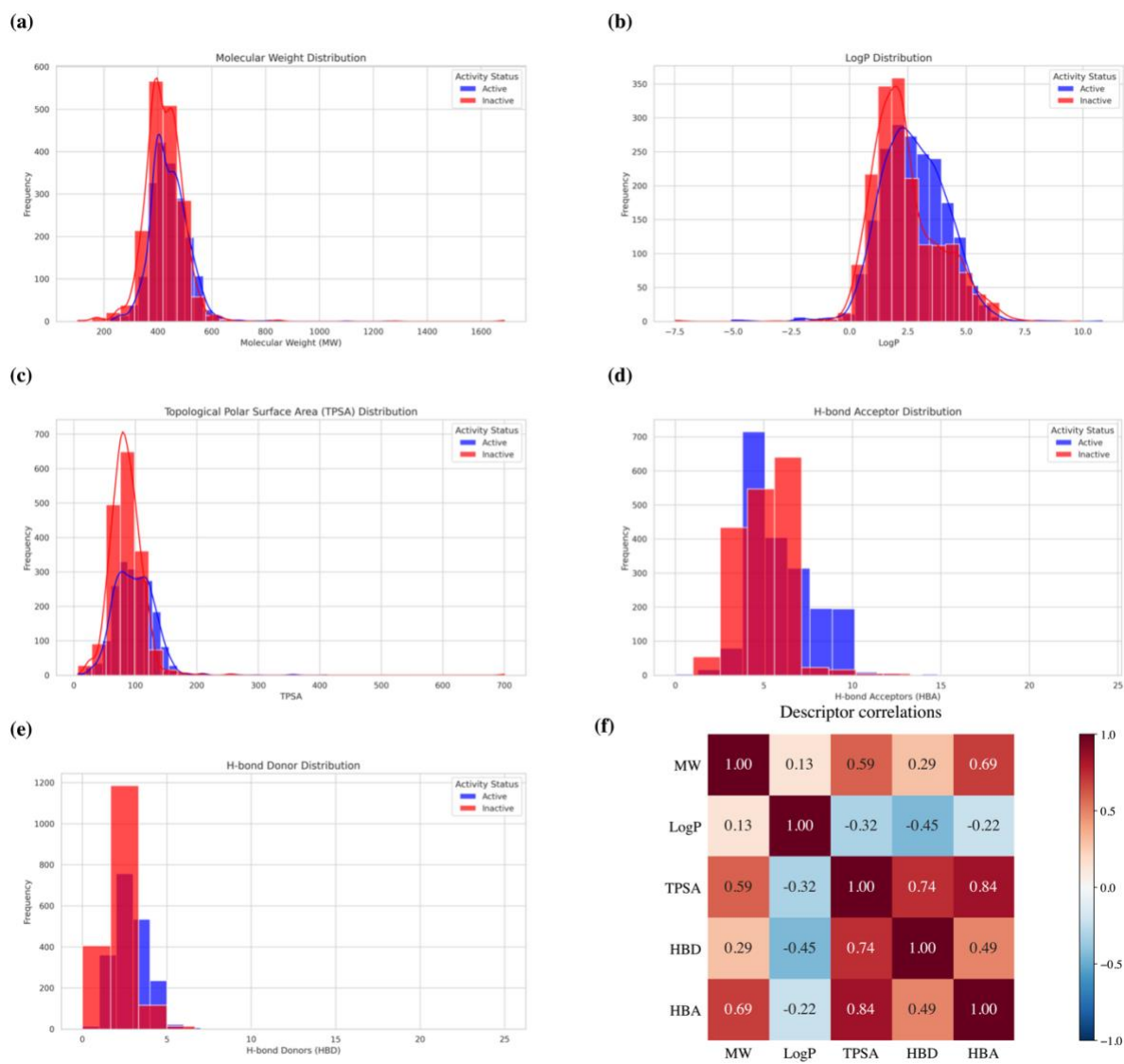

**Supplementary Figure 6. Panels (a–e):** original supplied plots. Blue = Active; red = source negative class (labelled “Inactive”). Panel (f): coefficients transcribed from the supplied heatmap, displayed on a symmetric scale centred at zero.

**Supplementary Table 1.** Overall alteration frequency.

| **Cancer Type** | **Alteration** | **Frequency** |
| --- | --- | --- |
| Skin Cutaneous Melanoma | Mutation | 4.3% |
|  | Amplification | 0.23% |
| Uterine Corpus Endometrial Carcinoma | Mutation | 3.78% |
|  | Amplification | 0.57% |
| Bladder Urothelial Carcinoma | Mutation | 0.93% |
|  | Amplification | 2.19% |
|  | Deep Deletion | 0.73% |
| Ovarian Serous Cystadenocarcinoma | Mutation | 0.68% |
|  | Amplification | 2.05% |
|  | Deep Deletion | 1.03% |

**Supplementary Table 2. Dataset partitioning for QSAR machine learning development.** Summary of active and decoy compound distributions across training (optimization) and test (external validation) sets.

| **Dataset** | **Number of Compounds** | **Percentage** |
| --- | --- | --- |
| Training set | 5,524 | 80% |
| Test set | 1,381 | 20% |
| Total | 6,905 | 100% |

**Supplementary Table 3.** Distribution of active and inactive molecular compounds in training and test datasets.

| **Dataset** | **Class** | **Number of Compounds** | **Percentage (%)** |
| --- | --- | --- | --- |
| Training | Active | 2,936 | 53.15 |
| Training | Inactive | 2,588 | 46.85 |
| Test | Active | 734 | 53.15 |
| Test | Inactive | 647 | 46.85 |
| Total | Active | 3,670 | 53.15 |
| Total | Inactive | 3,235 | 46.85 |

**Supplementary Table 4.** Binding affinity (DG) scores of the top-ranked phytocompounds against 4X61

| Compound ID: | 2D Structure | Docking score (kcal mol<sup>-1</sup>) |
| --- | --- | --- |
| CID 44345182 | 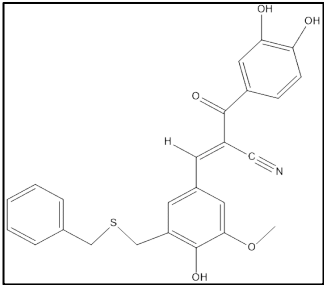 | -9.8 |
| CID 2794188 | 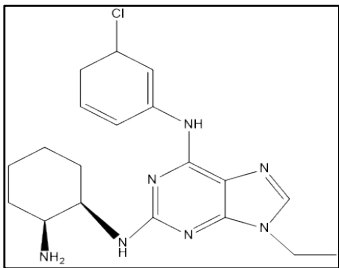 | -9.7 |
| CID 44345334 | 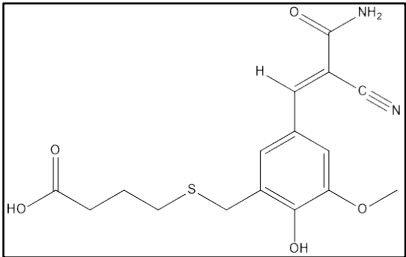 | -7.8 |
| CID 90241673 (control) | 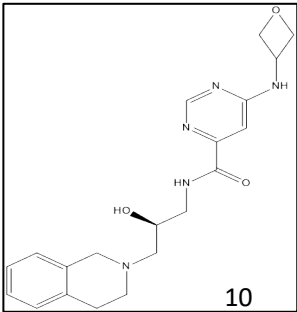 | -9.2 |
